# Exploring the Self-Assembly of Encapsulin Protein Nanocages from Different Structural Classes

**DOI:** 10.1101/2021.06.06.447285

**Authors:** India Boyton, Sophia C. Goodchild, Dennis Diaz, Aaron Elbourne, Lyndsey Collins-Praino, Andrew Care

## Abstract

Encapsulins, self-assembling icosahedral protein nanocages derived from prokaryotes, represent a versatile set of tools for nanobiotechnology. However, a comprehensive understanding of the mechanisms underlying encapsulin self-assembly, disassembly, and reassembly is lacking. Here, we characterise the disassembly/reassembly properties of three encapsulin nanocages that possess different structural architectures: *T* = 1 (24 nm), *T* = 3 (32 nm), and *T* = 4 (42 nm). Using spectroscopic techniques and electron microscopy, encapsulin architectures were found to exhibit varying sensitivities to the denaturant guanidine hydrochloride (GuHCl), extreme pH, and elevated temperature. While all encapsulins showed the capacity to reassemble following GuHCl-induced disassembly (within 75 min), only the smallest *T* = 1 nanocage reassembled after disassembly in basic pH (within 15 min). Furthermore, atomic force microscopy revealed that all encapsulins showed a significant loss of structural integrity after undergoing sequential disassembly/reassembly steps. These findings provide insights into encapsulins’ disassembly/reassembly dynamics, thus informing their future design, modification, and application.

## INTRODUCTION

Protein nanocages (e.g., virus-like particles (VLPs), ferritins, heat-shock proteins) self-assemble from multiple protein subunits into highly-organised macromolecular structures, that exhibit well-defined inner cavities, outer surfaces, and interfaces between subunits. Their capacity to encapsulate cargo, coupled with the ability to genetically and/or chemically modify their structures, has enabled protein nanocages to be custom-engineered for a multitude of applications, including biocatalysis, materials synthesis, sensing, vaccines, and drug delivery.^1, 2^

Encapsulins are an emerging class of protein nanocages found inside many archaea and bacteria. They self-assemble from identical protein subunits into hollow icosahedral nanocages that structurally resemble the major capsid protein gp5 of the HK97 virus.^3, 4^ Based on their triangulation number (*T*), all encapsulins exhibit one of the following three symmetrical icosahedral architectures: *T* = 1 (24 nm, 60-mer, 12 pentameric units); *T* = 3 (32 nm, 180-mer, 12 pentameric and 20 hexameric units); and *T* = 4 (42 nm, 240-mer, 12 pentameric and 30 hexameric units). ^5–8^ In nature, encapsulins house cargo enzymes that mediate oxidative stress resistance, iron storage, anaerobic ammonium oxidation, or sulfur metabolism.^9–11^ Uniquely, encapsulins selectively self-assemble around cargo enzymes tagged with a small encapsulation signal peptide (ESig), packaging them.^12^ This mechanism has been adapted to load foreign cargo into encapsulins, reprogramming their functionality for different practical applications. ^13–16^

Encapsulin subunits autonomously assemble, with extraordinary fidelity, into macromolecular nanocages. Such self-assembly is not only driven by folding of the individual polypeptide chains, but also by dynamic noncovalent interactions between the different polypeptide chains both within subunits and at the interfaces between subunits in the assembled supramolecular structure.^17^ Unravelling the self-assembly mechanisms of protein nanocages is complicated, especially if they exhibit highly symmetric homooligomeric structures, like encapsulins.^18^ Nevertheless, multiple analytical techniques now allow the molecular mechanisms underlying protein nanocage assembly (e.g., protein folding) to be characterised and subsequently exploited.

For instance, the disassembly/reassembly of protein nanocages belonging to the ferritin family have been studied via a combination of: intrinsic tryptophan fluorescence (ITF), circular dichroism (CD), UV/VIS spectroscopy and synchrotron small-angle X-ray scattering (SAXS) measurements to assess protein conformation^19–22^, transmission electron microscopy (TEM) and dynamic light scattering (DLS) to evaluate structural integrity, shape and size distribution, and laser light scattering to monitor assembly kinetics.^23^ One study revealed that ferritin disassembles at extremely acidic pH 1.5, then shows a rapid reassembly upon return to neutral pH 7.0, accompanied by folding, followed by a slow phase in which the final 24-mer nanocage is formed.^23^ Importantly, this fundamental work led to the rational re-design of ferritin subunit interfaces, resulting in engineered nanocages capable of disassembly at a more amenable pH 4.0.^24, 25^ Such modification now permits labile compounds (e.g. small-molecule drugs) to be controllably loaded into ferritin nanocages in a facile and non-destructive manner, enabling downstream applications (e.g., drug delivery).^24, 25^

In contrast, experimental data pertaining to encapsulins’ ability to disassemble/reassemble and the mechanisms that underpin this natural phenomenon are sparse. The most characterised system is the *T* = 1 encapsulin from *Thermotoga maritima* (*Tm-Enc*), whose disassembly/reassembly has been primarily inspected via CD, polyacrylamide gel electrophoresis (PAGE), and TEM.^14^ Specifically, *Tm-Enc* has been found to disassociate in strong acidic and alkaline conditions, or high concentrations of denaturing agents (e.g., guanidine hydrochloride, GuHCl). Furthermore, *Tm-Enc* was shown to spontaneously reassemble upon returning to the initial conditions (i.e., neutral pH or absence of denaturant).^27^ Interestingly, *Tm-Enc* can be reassembled in the presence of ESig-tagged cargo (e.g., proteins, nanomaterials), resulting in their selective encapsulation *in vitro*, and thus further expanding encapsulins’ utility.^8, 27, 28 29^ Despite these promising findings, key questions concerning the biophysical mechanisms and physicochemical factors that underlie encapsulin disassembly/reassembly, and how they might be controlled, remain unanswered, especially for the *T* = 3 and *T* = 4 nanocages.

Motivated by this absence of information, we selected encapsulins with structures representing each of the three known architectures, and then interrogated their disassembly/reassembly. These nanocages included *Tm-Enc* (*T* = 1), and the larger and more structurally complex encapsulins from *Myxococcus xanthus* (*Mx-Enc*, *T* = 3) and *Quasibacillus thermotolerans* (*Qt-Enc, T* = 4). We combined intrinsic tryptophan fluorescence (ITF) spectroscopy, DLS, PAGE and TEM, to accurately monitor the assembly states of all three encapsulins under varying physicochemical conditions, including exposure to extreme pH, strong denaturants (GuHCl), and elevated temperatures. Furthermore, the effect disassembly/reassembly had on the nanocages’ structural integrity was evaluated by atomic force microscopy (AFM). Together, this work provides critical insights into the dynamic mechanisms that govern the disassembly/reassembly of differing encapsulin structures, which will help to expedite and broaden their future design, modification, and practical application.

## MATERIALS AND METHODS

### Materials

All chemicals and reagents used in this study were purchased from Sigma-Aldrich, unless stated otherwise.

### Molecular cloning of constructs

All inserts were codon optimised for expression in *Escherichia coli* and custom synthesised as gBlock Gene Fragments (Integrated DNA Technologies). Encapsulins from *Thermotoga maritima (Tm)* (UniProt: TM_0785), *Myxococcus xanthus (Mx)* (UniProt: MXAN_3556) and *Quasibacillus thermotolerans (Qt)* (UniProt: QY95_01592) were each synthesised with flanking restriction sites (NcoI/BamHI). For gene expression in *E. coli*, *Tm-Enc* was cloned into pETDuet-1 (Novagen, Merck), and *Mx-Enc* and *Qt-Enc* were cloned into pACYC-Duet-1 (Novagen, Merck), summarised in Supplementary Table S1. *E. coli* α-Select (Bioline, UK) was used for general plasmid storage and propagation. Gene insertion was confirmed by PCR using primer pairs pETUpstream/DuetDOWN (Merk). *E. coli* BL21 (DE3) cells (New England Biolabs) were used for recombinant protein expression. Herein, cells were transformed with the appropriate plasmids, and the resulting transformants were selected on Luria–Bertani (LB) agar supplemented with either 100 mg/mL of carbenicillin or 50 mg/mL of chloramphenicol (see Supplementary Table S1).

### Recombinant Protein Production

Protein expression experiments were performed in LB medium supplemented with carbenicillin (100 mg/mL) or chloramphenicol (50 mg/mL). Briefly, strains were streaked out on LB agar plates and grown overnight at 37°C. A starter culture (1 colony in 5 mL LB) was grown for 16 h at 37°C and used to inoculate 500 mL of LB media. Cultures were incubated at 37°C/200–250 rpm until an optical density at 600 nm (OD600) of 0.5-0.6 was reached. Protein synthesis was then induced by the addition of isopropyl-β-d-thiogalactopyranoside (IPTG) to a final concentration of 0.1 mM. Induced cultures were incubated at 37°C/200–250 rpm for 4 h and then cells were harvested via centrifugation (8,000 × *g*, 4°C, 15 min). The resulting cell pellets were stored at −30°C until further use.

### Protein purification

Cell pellets from 500 mL encapsulin-producing cultures were thawed and resuspended in 30 mL of lysis buffer (50 mM 4-(2-hydroxyethyl)-1-piperazineethanesulfonic acid (HEPES) buffer pH 7.4 (Chem-Supply Pty) and Benzonase® nuclease 10 U/mL). Cells were lysed by three rounds of passage through a French pressure cell at 1000 psi and centrifuged at 8,000 × *g*, 4°C for 15 min. Supernatant containing soluble protein was heat treated in a water bath at 65°C for 15 min before centrifugation (10,000 × *g*, 4°C, 10 min). Protein precipitation was initiated by adding 10 % (w/v) PEG8000 and 2 % (w/v) NaCl to the supernatant, followed by incubation on ice for 30 min. Next, the sample was spun down at 10,000 × *g* for 10 min at 4°C. The precipitated protein was resuspended in 2.5 mL of HEPES buffer (50 mM, pH 7.4) and filtered through a 0.22 mm syringe filter.

All purifications were carried out on an Äkta™ start chromatography system (GE Healthcare). The three encapsulins used in this study were purified via size exclusion chromatography (SEC) using a HiPrep 26/60 Sephacryl S-500 HR column (GE Healthcare) equilibrated with 50 mM HEPES pH 7.4. Fractions showing the corresponding encapsulin band on SDS-PAGE were pooled and subjected to further purification via anion-exchange chromatography using a HiPrep Q 16/10 column (GE Healthcare) equilibrated with 50 mM HEPES pH 7.4. Encapsulin proteins were eluted with linear gradient of 0–0.3 M NaCl and 0.3– 1M NaCl (Supplementary Figure S1). Fractions containing encapsulin, identified via sodium dodecyl sulphate polyacrylamide gel electrophoresis (SDS-PAGE), were pooled, concentrated and buffer exchanged into 50 mM HEPES buffer pH 7.4 using Amicon Ultra-15 centrifugal filter units with a 100 KDa cut-off. Lastly, purified encapsulin Enc proteins were filtered through a 0.22 mm syringe filter and stored at −30°C until further use.

### PAGE analysis and protein quantification

Protein samples were denatured, separated, and visualised using SDS-PAGE, with molecular weights compared with a commercial protein ladder (Precision Plus Protein, BioRad). The Bio-Rad mini-protean system (Bio-Rad laboratories) was used for SDS-PAGE analysis. Protein samples were mixed in a 1:1 ratio with 2X Laemmli sample buffer with 50 mM 1,4-dithiothreitol and heated at 99°C for 10 min at 300 rpm. Electrophoresis was performed at 200 V for 30 min on a 4–20% polyacrylamide gel (Mini-PROTEAN® TGX™, BioRad) in SDS running buffer (25 mM Tris, 192 mM glycine, 1% (w/v) SDS, pH 8.3). Gels were stained following the Coomassie G-250 safe stain protocol. Encapsulin assembly was visualised via non-denaturing Blue Native-PAGE (BN-PAGE). The XCell SureLock Mini-Cell Electrophoresis System (ThermoFisher Scientific) was used for BN-PAGE analysis. Protein samples were mixed in a 1:4 ratio with 4X Native-PAGE sample buffer (ThermoFisher Scientific) and loaded into NativePAGE™ 3–12% Bis-Tris protein gels (ThermoFisher Scientific). BN-PAGE was performed using two different running buffers: 1X anode buffer (NativePAGE™ running buffer 20X, ThermoFisher Scientific) in the outer buffer chamber and 1X dark blue cathode buffer (1X anode buffer, 0.02% (w/v) Coomassie G-250) in the inner buffer chamber. Lastly, the samples were run on ice at 150 V for 90 min follow by a second run at 250 V for 30 min. Protein concentration was determined by measuring the absorbance at 280 nm on a NanoDrop 2000 Spectrophotometer instrument (ThermoFisher Scientific).

### In vitro disassembly/reassembly of encapsulin

To characterise the *in vitro* disassembly of encapsulin, the presence of a denaturing agent (GuHCl), changes in pH, and thermally induced disassembly methods were used. GuHCl was added to the encapsulin sample to final concentrations between 1–7 M. For pH disassembly, dilution of the purified encapsulins in 50 mM HEPES at different pH (3–13) was performed. Final pH was confirmed with pH strips. Once in their respective conditions, encapsulin samples were incubated for 1 h at room temperature. For thermally induced disassembly, the encapsulin sample was subjected to a temperature ramp from 20–90°C at a rate of 2°C/min. In all disassembly methods, the final encapsulin concentration was 5 μM, dithiothreitol (DTT) was added to the encapsulin sample to a final concentration of 1 mM and 50 mM HEPES pH 7.4 was used to reach the desired final volume.

For stability experiments, 5 μM of encapsulin was incubated for 1 h at room temperature under reducing (10 mM DTT), oxidising (10 mM H2O2), and high ionic strength (1 M NaCl) conditions.

The subsequent reassembly of encapsulins was initiated by returning the sample to original conditions. Briefly, the samples were dialysed against 50 mM HEPES buffer pH 7.4 and 1 mM DTT at room temperature overnight, using 3.5K MWCO SnakeSkin dialysis tubing (ThermoFisher Scientific). To measure the reassembly rate, samples were removed from dialysis every 15 min for 75 min and centrifuged for 5 min at 10,000 × *g* to remove any aggregated proteins.

### Intrinsic tryptophan fluorescence

ITF measurements of encapsulins in their varying states of assembly were performed with an FP-8500 Spectrofluorometer (JASCO) using a 3 mm pathlength micro-volume quartz cuvette. Samples (60 μL) were prepared in triplicate with a final encapsulin concentration of 5 μM. Samples were excited at 290 nm and emission spectra were collected from 300–450 nm. The measurement parameters were: excitation and emission bandwidths of 5 nm, response of 0.2 s, medium sensitivity, data interval of 0.1 nm, scan speed of 100 nm/min, and 4 measurement accumulations were averaged. To investigate the effect of temperature on the different encapsulins, spectra were collected from 20–90°C with a temperature ramp of 2°C/min. The obtained spectra were further processed by buffer spectra subtraction using Spectra Manager™ software (JASCO), and the ratio between the fluorescence intensity at 360 and 330 nm (360/330) was calculated and plotted in Microsoft Excel.

### Transmission electron microscopy (TEM)

To visualise morphology, size, and state of encapsulin assembly, TEM was performed using a Philips CM10 microscope (100 kV accelerating voltage). Briefly, encapsulin samples (0.2 mg/mL) were adsorbed onto pioloform-coated 200 mesh copper grids (ProSciTech) for 2 min. Prior to imaging, samples were negatively stained for 1 h using uranyl acetate replacement (UAR-EMS, Electron Microscopy Sciences), washed with ultrapure water, and allowed to dry for at least 15 min.

### Dynamic light scattering

To measure the diameter of encapsulins, DLS was performed with a Malvern Zetasizer ZSP instrument equipped with a 633 nm laser. Samples with a final encapsulin concentration of 5 μM were prepared as described above. Three measurements were performed at 25°C, using a plastic microcuvette (ZEN0040, Malvern), 173° backscatter and automatic attenuator selection. Data analysis was performed in the Zetasizer Nano software. All DLS sizes reported, herein, are size by number values calculated using distribution analysis. A 1 cm quartz cuvette was used for temperature ramp experiments. Heat maps were made using GraphPad Prism 9.

### Atomic Force Microscopy (AFM)

Images and force curve measurements were obtained using a Cypher ES Atomic Force Microscope (Oxford Instrument, Asylum Research, Santa Barbara, CA, USA) at room temperature (25°C). Protocols were adapted from Collett *et al* ^26, 27^. For imaging, the instrument was operated in amplitude modulated-AFM (AM-AFM), while force measurements were obtained in contact mode. Biolever BL-AC40TS cantilevers (Oxford Instruments, Asylum Research, Santa Barbara, CA, USA, nominal spring constant k_c_ = 0.09 N/m) were used for all measurements. All experiments were completed within a droplet of media (~100 μL) with a concentration of ~100 ng of encapsulin deposited onto freshly cleaved muscovite surface (the supporting substrate). Prior to experimentation, each cantilever was calibrated via the thermal spectrum method and the lever sensitivity was determined using force spectroscopy. Processing of AFM data involved using a combination of the Asylum Research software, custom MATLAB codes, and the Gwyddion software package ^28^.

### Determining elasticity of intact encapsulins

Force versus distance curves (FD curves) were obtained from the central region of encapsulins. Following image location, FD curves were first obtained on an area of bare mica next to the nanocage to ensure linear (non-elastic) FD curves were observed, which provided a reference for determining elasticity of the particles. The spring constant (k_c_) of the cantilever was determined for each cantilever used as described above (values in the range of 0.05–0.1 N/m were obtained for all cantilevers). The tip was then directed to the central region of the particle to obtain accurate indentation data. Upward of 50 FD curves were recorded across several particles for each sample.

The tip-sample contact point between the AFM cantilever and the encapsulin was performed independently for each FD curve analysed using methods previously described ^29^. Specifically, the contact point between the AFM cantilever tip and the encapsulin is defined as the point at which the cantilever first makes physical contact with the surface adsorbed particle. Following this point, the cantilever bends in response to interactions with the encapsulin, while the nanocage itself is indented. In a raw FD curve (z-displacement (Z) vs. cantilever deflection (d)), this point of first contact is mathematically defined as Z_0_ and d_0_, respectively. This is observed in the AFM force curve as an increase in force (y-axis), above the zeroed baseline, as a function of distance (x-axis prior to zeroing the indentation).

The indentation (δ) is then calculated via:

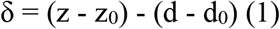

and transformed into indentation vs. force curves using Hooke’s law;

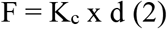

The elastic response of the samples is then fit to the Hertz/Sneddon equation (3) to obtain a Young’s elastic modulus (*E*) for each of the fitted curves. Using custom MATLAB scripts, curves were fitted to the equations:

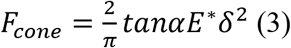

and;

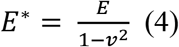

where F is loading force, δ is indentation depth, α is the cone opening angle, E* is the apparent elastic modulus, and v is the Poisson’s ratio. A conal tip angle of 34.4° and a Poisson’s ratio of 0.5 was used for processing of all force curves. All elasticity values were obtained from curves fitted with an R^2^ value of 0.9 or above. Data falling below this quality was rejected for further analysis.

## RESULTS AND DISCUSSION

### Monitoring encapsulin assembly/disassembly using intrinsic tryptophan fluorescence

Unloaded *Tm-Enc, Mx-Enc,* and *Qt-Enc* were produced in *E. coli* and purified by SEC and anion exchange chromatography prior to biophysical characterisation. Purification and correct self-assembly were confirmed using SDS-PAGE, blue native-PAGE, DLS, and TEM (Figure 2g, Supplementary Figure S1). TEM images of self-assembled *Tm-Enc, Mx-Enc,* and *Qt-Enc* displayed structures with consistent shape and size (Figure 2g) and DLS analysis indicated a diameter of 23.7 ± 4.9 nm for *Tm-Enc*, consistent with its crystal structure data^12^, and a diameter of 32.0 ± 6.0 nm for *Mx-Enc* and 38.2 ± 7.3 nm for *Qt-Enc,* consistent with their Cryo-EM structures.^6, 8^ *Mx*-*Enc* was produced primarily in its *T* = 3 structure; however, populations of smaller, *T* = 1 like, structures were evident in Native-PAGE and TEM results (~18 nm) (Supplementary Figure S1, Figure 2g). This variation in size of *Mx-Enc* has been previously observed, where recombinantly produced *Mx-Enc* without the presence of ESig-tagged cargo will form heterogenous *T* = 1 and *T* = 3 populations.^8^

Tryptophan fluorescence is extremely sensitive to local environment. As such, protein unfolding, disassembly or conformational transitions often result in a change in emission spectra of the Trp(s) within a protein; a lower maximum wavelength (blue-shifted) when the Trp(s) are buried, and a higher maximum wavelength (red-shifted) when solvent exposed.^30^

Encapsulin subunits adopt a HK97-fold and have three conserved structural regions, a peripheral (P)-domain, axial (A)-domain, and an elongated (E)-loop region (Supplementary Figure S2).^3, 4^ Conveniently, each subunit of *Tm-Enc, Mx-Enc,* and *Qt-Enc* contains five, three, and two Trp residues, respectively. For each encapsulin, at least one of these Trps is located within the interface between subunits, and one is located within the hydrophobic core of a single subunit (Figure 1a, b & c). Therefore, these intrinsic Trp residues are likely to be suitable reporters for both assembly/disassembly of the encapsulin macrostructure and folding of the individual subunit polypeptide chains. Indeed, intrinsic Trp fluorescence (ITF) has previously been used to monitor the re-folding of *T* = 1 encapsulin from *Rhodococcus erythropolis* N771 when desorbed from a zeolite substrate.^31^ ITF is also an appealing technique to monitor the process of encapsulin assembly/disassembly owing to its relative simplicity, ability to report on a dynamic ensemble of structures in solution and non-destructive nature, which enables measurements to be performed in real-time. Additionally, unlike fluorescence resonance energy transfer (FRET)-based techniques, ITF does not require modification of the protein with any extrinsic labels that may alter the assembly/disassembly dynamics. Although circular dichroism (CD) may provide more specific secondary structural information, unlike ITF, CD is not compatible with the high concentration of GuHCl and extreme pH likely required for encapsulin disassembly.^32^

**Figure 1.**
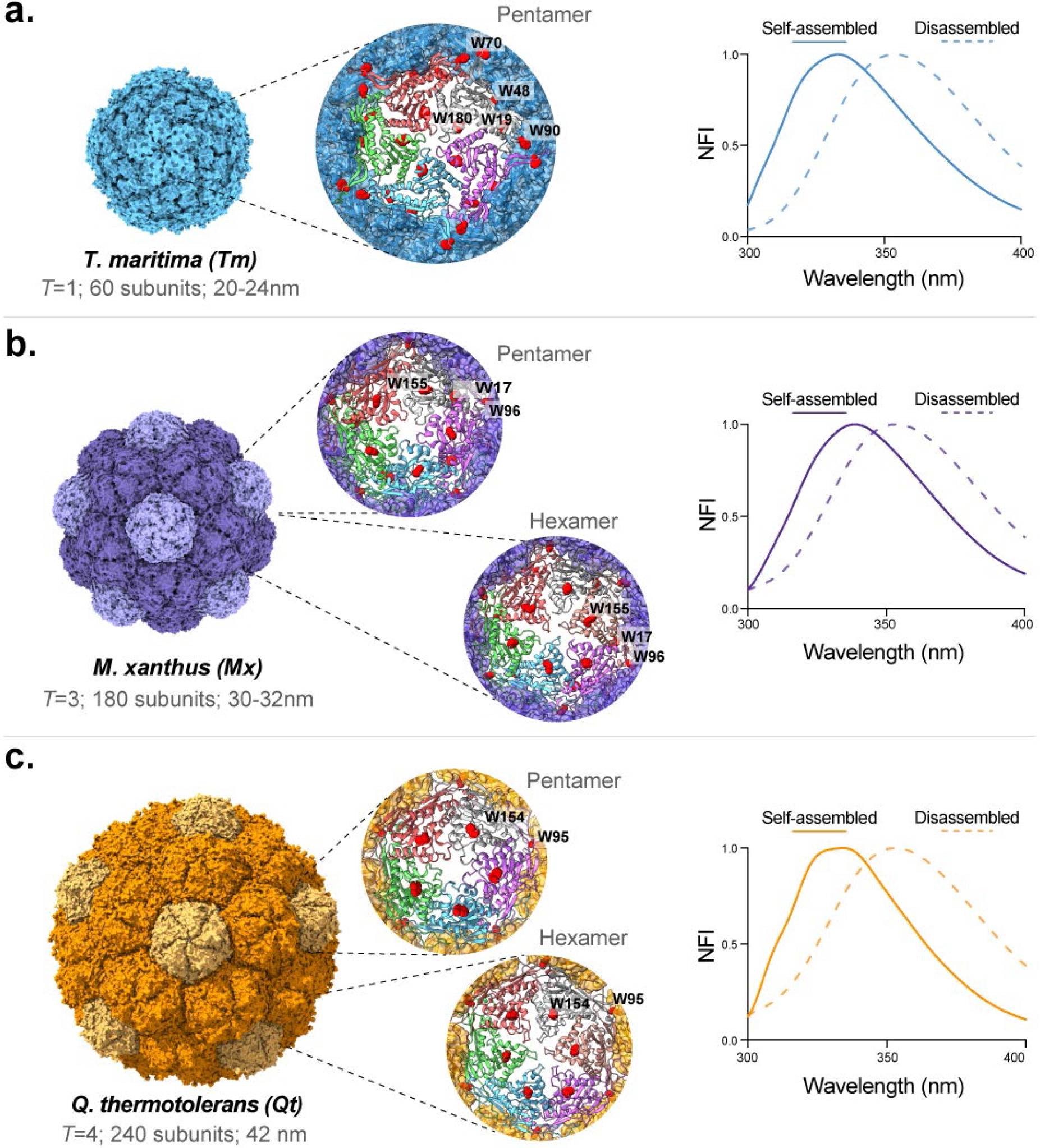
Monitoring encapsulin nanocage assembly states via ITF. Schematic diagram showing the assembled architectures of **(a)** *Tm-Enc* (PDB: 3DKT), **(b)** *Mx-Enc* (PDB: 4PT2), and **(c)** *Qt-Enc* (PDB: 6NJ8). For each structure, the pentameric and hexameric units are shown in light and saturated colours, respectively. Within the expanded pentameric or hexameric units, the tryptophan (W) residues belonging to individual subunits are highlighted in red: *Tm*: W19, W48, W70, W90 and W180; *Mx*: W17, W96 and W155; *Qt*: W95 and W154. Molecular graphics were created using UCSF ChimeraX ^36^. Using ITF, a shift in emission discriminated between the maximum emission wavelength of buried Trp in ‘assembled’ form: ~334 nm for **(a)** *Tm-Enc* and **(c)** *Qt-Enc,* or ~338 for **(b)** *Mx-Enc*; and solvent exposed Trp in ‘disassembled’ form (~354 nm) upon addition of 7 M GuHCl. Emission wavelengths are represented as Normalised Fluorescence Intensity (NFI).

Figure 1 shows the ITF of assembled *Tm-Enc*, *Mx-Enc* and *Qt-Enc* overlayed with the spectra obtained after 1 h incubation with 7 M GuHCl. For all three assembled encapsulins, blue-shifted Trp fluorescence spectra are observed (~334 nm maximum emission wavelength for *Tm-Enc* and *Qt-Enc;* and ~338 nm for *Mx-Enc*), while in the presence of 7 M GuHCl, a dramatic red-shift of the fluorescence spectra is seen (~354 nm maximum emission wavelength). A maximum emission wavelength over 350 nm is observed where the Trps within a protein are completely solvent exposed, as in an unfolded polypeptide chain.^33^ Therefore, it appears that 7 M GuHCl fully disassembles *Tm-Enc*, as has previously been demonstrated^14^, as well as both *Mx-Enc* and *Qt-Enc,* by completely unfolding the protein.

It is also interesting to note that the overall fluorescence emission spectral shape represents an average of all Trp environments present in the protein.^34^ Where a protein contains multiple Trps, these may exist in different environments and therefore emit at different wavelengths. While we cannot draw conclusions about the foldedness/assembly of *Tm-Enc*, *Mx-Enc* and *Qt-Enc* relative to one another (as they all contain a different number of Trps), changes in the spectral shape for the same encapsulin may discern encapsulin assembly (i.e., interaction between subunits) as opposed to the folding of individual subunit polypeptide chains. This is particularly true for *Qt-Enc*. As can be seen in Figure 1f, the peak-shape for assembled *Qt-Enc* is flattened. This is consistent with its two Trp residues being located in different solvation environments such that the spectra two Trps overlap, but are not resolved (i.e., Trp95 being more buried in a helix at the pentamer/hexamer interface than Trp154, which is located on a loop in the A-domain, nestled within the hydrophobic core of a single *Qt-Enc* subunit) (Figure 1c, Supplementary Figure S2).

Instead of reporting the maximum fluorescence wavelength, herein, we have presented much of our ITF data as a fluorescence intensity ratio of emission at 360 nm and 320 nm (360/320). This ratio is used as a proxy for the shift in overall Trp fluorescence emission peak, and avoids any bias arising from the spectral peak shape.^35^ This approach has allowed us to expand the repertoire of methods used to monitor encapsulin assembly/disassembly across a range of conditions.

### GuHCl induced encapsulin disassembly

GuHCl is a denaturant that affects protein structure by disrupting non-covalent interactions, including hydrogen bonding and hydrophobic effects.^37^ Our initial ITF results indicated that 7 M GuHCl completely unfolds the *Tm-Enc*, *Qt-Enc* and *Mx-Enc* protein polypeptide chains (Figure 1), and thus, disassembles the encapsulin macrostructure. To further examine the effect of GuHCl on encapsulin folding and assembly, all encapsulins were incubated in varying concentrations of GuHCl (0–7 M) for 1 h before performing ITF and DLS.

The observed shift in Trp emission peak for *Tm-Enc*, *Mx-Enc* and *Qt-Enc* in the presence of GuHCl are shown in Figure 2a. As expected from our previous results, all three encapsulins show a blue to red-shift from the assembled (0 M GuHCl, Δ360/320 = 0) to unfolded state (7 M GuHCl, Δ360/320 = 1). However, the concentration of GuHCl at which this transition occurs varies significantly between the different encapsulins. *Tm-Enc* Trp solvation began from approximately 3 M GuHCl, with complete Trp solvation from 6 M GuHCl, while *Mx-Enc* and *Qt-Enc* Trp solvation began at a lower concentration of 1 M GuHCl, with complete Trp solvation from 4 M and 5 M GuHCl, respectively. This variation may reflect structural complexity, with the smaller and pentameric *Tm-Enc* being the most robust to GuHCl compared to the larger pentameric and hexameric *Mx-Enc* and *Qt-Enc*.

**Figure 2.**
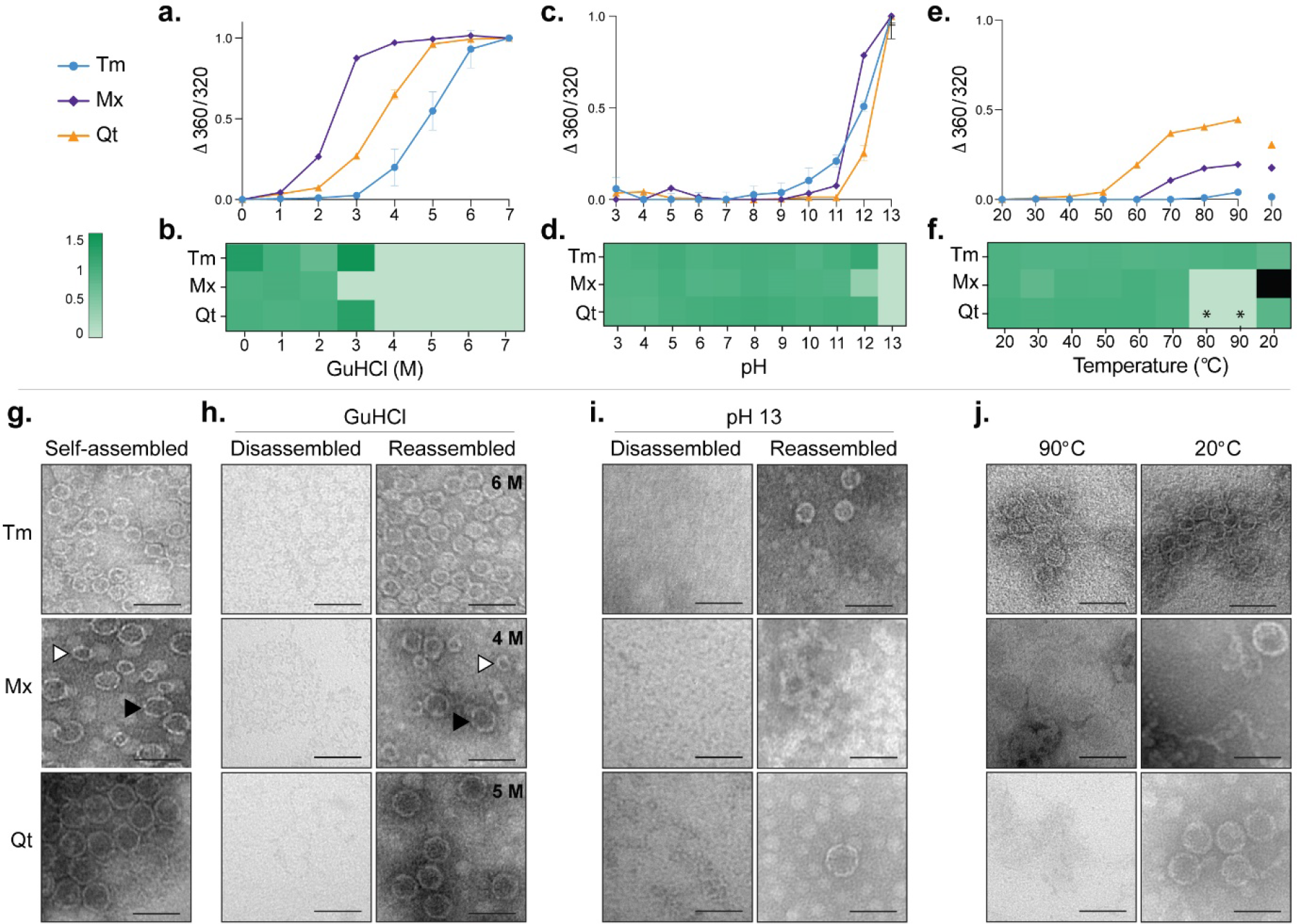
GuHCl, pH, and thermally induced disassembly of Encs. **(a)** ITF showed an observed shift in emission wavelength (360/320) and indicated Trp solvation begins from 1 M GuHCl for *Mx-Enc* and *Qt-Enc*, and 3 M for *Tm-Enc*. **(b)** DLS measurements indicated *Mx-Enc* disassembled into its subunits (<0.5 nm) from 3 M GuHCl, and *Tm-Enc* and *Qt-Enc* disassembled from 4 M GuHCl. **(c)** ITF showed Trp solvation only significantly increased in alkaline conditions from pH 12 for *Tm-Enc, Mx-Enc* and *Qt-Enc*. **(d)** DLS measurements indicated *Mx-Enc* began to disassemble at pH 12 (~13 nm) and all Enc disassembled into their subunits at pH 13 (<0.7 nm). **(e)** ITF emission wavelengths (360/320) of Enc between 20–90°C and cooled back to 20°C. *Tm-Enc* remained stable with slight Trp solvation from 80°C that reversed when cooled back to 20°C. *Mx-Enc* began to display Trp solvation from 60°C, and *Qt-Enc* from 40°C. **(f)** DLS measurements show *Tm-Enc* remained assembled to 90°C, and *Mx-Enc* and *Qt-Enc* began to disassemble into their subunits (<0.5 nm) at 80°C. *A smaller population of intermediate *Qt-Enc* structures remained at 80-90°C. Upon cooling to 90°C, *Qt-Enc* appeared to reassemble, whereas *Mx-Enc* aggregated (indicated by black square). **(g-j)** TEM images show self-assembled encapsulins compared to disassembled encapsulins under varying conditions and subsequent reassembly back into spherical nanocages; Mx-Enc (black triangle *T*=3, white triangle *T*=1) (scale bars = 50 nm). For ITF data, the difference in emission wavelength of complete Trp solvation (normalised to 1) and assembled Enc (normalised to 0) was plotted. Error bars represent the mean ± standard deviation; n = 3 from three independent experiments. DLS results were normalised so that 1 = expected assembled size and 0 = disassembled encapsulin.

To complement these ITF results, DLS analysis was performed to characterise the size distribution of encapsulin in solution for the same samples (Figure 2b, Supplementary Table S2). For *Mx-Enc*, no intact encapsulin macrostructure is detectable by DLS at concentrations ≥ 3 M GuHCl, which also correlates with the significant red-shifted Trp emission at 3 M GuHCl (Δ360/320 = 0.88). This result suggests that *Mx-Enc* is both disassembled and largely unfolded at GuHCl concentrations ≥ 3 M. However, for *Tm-Enc* and *Qt-Enc*, absence of the intact encapsulin macrostructure occurs at lower GuHCl concentrations than the major red-shift in Trp emission. For example, at 4 M GuHCl, the *Tm-Enc* macrostructure is not detectable by DLS, but the Trp emission remains significantly blue-shifted (Δ360/320 = 0.20), corresponding to a native-like fold. A similar result is seen for *Qt-Enc* at 3 M GuHCl (Δ360/320 = 0.27). TEM and native-PAGE images of each encapsulin after incubation in the lowest GuHCl concentration required for significant Trp solvation also show the absence of nanocage macrostructures, confirming their disassembly. Upon dialysis in buffer (50 mM HEPES, 1 mM DTT) overnight, all encapsulins subsequently reassembled into their original macrostructure (Figure 2h, Supplementary Figure S3).

Taken together, these results point to the existence of an intermediate state(s) in *Tm-Enc* and *Qt-Enc* GuHCl-induced disassembly, in which the nanocage breaks down into smaller (<0.5 nm) entities with a native-like fold. There is insufficient evidence from ITF or DLS to define the stoichiometry or structure of this intermediate state(s). However, a previous native mass spectrometry study of the *T* = 1 encapsulin from *B. linens* suggested reassembly occurs via formation of stable dimers prior to the final nanocage.^38^ It is possible that disassembly of *Tm-Enc* and *Qt-Enc* proceeds via a similar mechanism. We also cannot rule out that *Mx-Enc* proceeds via an equivalent intermediate state upon GuHCl-induced disassembly. However, if this is the case, our data suggests the chemical stability of the nanocage and the intermediate for *Mx-Enc* are more closely matched, and less chemically stable, than those of *Tm-Enc* or *Qt-Enc,* such that both disassembly and complete unfolding occur at close to 3 M GuHCl. This potential model of encapsulin disassembly warrants further investigation, as disassembly of the nanocage utilising lower GuHCl concentrations, without completely unfolding the encapsulin protein, may provide an effective scheme for loading and/or releasing more chemically sensitive cargo.

Overall, our results for GuHCl-induced encapsulin disassembly suggest that folding plays a critical role for encapsulin nanocage assembly/disassembly. The smaller, less complex *Tm-Enc* structure requires more GuHCl for unfolding than the larger, more complex *Qt-Enc* and *Mx-Enc*. This agrees with the difference in the high stability of ferritin, which is a small protein nanocage comprised of 24 subunits, compared to the high sensitivity of larger and more structurally complex VLPs in denaturants. Ferritin is stable up to 6 M GuHCl^39^, and requires 8 M of denaturant urea for disassembly.^40^ A concentration of 2 M GuHCl is not strong enough to disassemble ferritin, but is able to unfold the α-helices surrounding ferritin channels, thereby enlarging its pores to allow entry of cargo without the need for complete disassembly.^41^ Contrastingly, the larger P22 VLP, which assembles from 420 subunits into a *T* = 7 nanoparticle-like structure that is 56 nm in diameter, dissociates at just 3 M GuHCl.^42^ However, the chemical stability of the encapsulins tested is not directly related to their size, as *Qt-Enc* requires more GuHCl for unfolding than the smaller *Mx-Enc*. Hence, other factors, such as the symmetry and stability of intermediate state(s), also needs to be taken into consideration.

### pH induced encapsulin disassembly

To investigate the role of electrostatic interactions in maintaining encapsulin macrostructure, ITF and DLS were also performed for the three encapsulins at varying pH (Figure 2c & d). The theoretical pI of *Tm-Enc*, *Mx-Enc* and *Qt-Enc* are 4.90, 5.45 and 5.02, respectively.^43^ If electrostatic interactions play a major role in the mechanism of encapsulin assembly, we would expect to see disassembly of the encapsulin nanocage at pH < pI (where the overall charge on the protein would be positive rather negative, as at neutral pH). However, this is not the case. ITF indicates that all encapsulins remain relatively stable across a broad pH range, with a significant change in Trp exposure only observed in extreme alkaline conditions (pH 12-13) (Figure 2c). DLS demonstrates the presence of assembled encapsulin at all pHs, with the exception of pH 13, and smaller structures (~13 nm) at pH 12 for *Mx-Enc* (Figure 2d, Supplementary Table S3). Disassembly of the nanocage structures at pH 13 was also confirmed by TEM (Figure 2i). Therefore, loss of electrostatic interactions between and/or within encapsulin subunits is unlikely to be a major driving force for disassembly. Disassembly more likely arises at extreme alkaline pH due to deprotonation of the protein side chains, resulting in a loss of the hydrogen bonding that holds the protein scaffold together.^44^

### Thermal encapsulin disassembly

To assess encapsulin thermostability, assembled samples were heated to 90°C with ITF and DLS measurements taken every 10°C up to 90°C, as well as upon return to 20°C. Additionally, encapsulins were heated to 90°C with TEM samples prepared both immediately after heating, and also after being cooled at 20°C for 1 h. ITF results showed *Tm-Enc* encapsulin remained stable throughout heating, with only a very slight observed red-shift in emission wavelength at temperatures above 80°C, which was recovered when cooled back to 20°C (Figure 2e). No significant change in size of the *Tm-Enc* nanocages was observed by DLS, and TEM images showed assembled cages after heating to 90°C, suggesting that the *Tm-Enc* nanocage is resistant to thermal disassembly (Figure 2f & j, Supplementary Table S4).

In contrast to *Tm-Enc*, both *Mx-Enc* and *Qt-Enc* disassemble at elevated temperatures. *Mx-Enc* Trp solvation began from 60°C and disassembled from 80°C, as shown by DLS (Figure 2e & f, Supplementary Table S4), and appeared disassembled at 90°C via TEM imaging (Figure 2j). Additionally, in contrast to *Tm-Enc*, disassembly of *Mx-Enc* by temperature is not reversible, as when cooled back to 20°C only large aggregates were found (Figure 2f, Supplementary Table S4). Similarly, ITF showed *Qt-Enc* began Trp solvation from 40°C and DLS indicated the majority of sample disassembled at 80°C and 90°C. However, even at 90°C, a smaller population (approx. 16%) of *Qt-Enc* subunits remained in intermediate structures of ~13 nm (Figure 2f, Supplementary Table S4). When cooled back to 20°C, DLS and TEM indicated *Qt-Enc* reassembled to its original size and ITF showed a blue-shift of emission (Figure 2e, f & j, Supplementary Table S4).

The difference in thermal stability and reassembly of *Tm-Enc*, *Mx-Enc* and *Qt-Enc* points to kinetic complexity in the encapsulin assembly pathway. For *Mx-Enc*, aggregation of the unfolded/disassembled state is likely the main obstacle to achieving proper reassembly. Our GuHCl data also suggest that the *Mx-Enc* unfolded/disassembled state(s) is less stable than for *Tm-Enc* or *Qt-Enc*. Whilst *Qt-Enc* began Trp solvation at 40°C, DLS showed it did not change in size until 80°C, indicating *Qt-Enc* undergoes structural change from 40°C, but remains assembled until 80°C. The stability of *Tm-Enc* and *Qt-Enc* at high temperatures are consistent with them being derived from thermophilic bacteria.^6, 12^ Whereas *Mx-Enc* is derived from bacteria that lives in soil^45^, which may explain its lower tolerance to temperature.

### Alternate disassembly conditions

We have begun to use the ITF technique, developed herein, to investigate other chemical additives that may alter the folding and/or assembly of encapsulins, including redox conditions (10 mM DTT or H_2_O_2_) and ionic environments (1 M NaCl). However, we are yet to find another condition to induce disassembly. All Encs remained stable in NaCl, and only *Mx-Enc* displayed a small blue-shift in emission in H_2_O_2_, and an observable increase in diameter after incubation in either DTT or H_2_O_2_ (Supplementary Figure S4, Supplementary Table S7). This may be due to reduction/oxidation of disulfide bonds causing a weaker association between subunits, resulting in swelling of the nanocage.^46^

### Encapsulin Reassembly

Based on the above results, high concentrations (6 M) of GuHCl and pH 13 conditions were selected for further analysis of the *Tm-Enc*, *Mx-Enc*, and *Qt-Enc* reassembly mechanism. Notably, these conditions disassemble encapsulin to differing degrees. ITF revealed maximum peak emissions for disassembly with pH 13 to be ~341 nm for *Tm-Enc*, and ~347 nm for *Mx-Enc* and *Qt-Enc*, whereas disassembly with 6 M GuHCl resulted in a maximum peak emission at ~354 nm for all encapsulins (Figure 3a–c). As previously discussed, a Trp emission maxima ≥350 nm (as observed for all encapsulins in 6 M GuHCl) is expected for unfolded protein. However, in pH 13, all encapsulins display a Trp max emission < 350 nm, suggesting that at least some of their subunits’ secondary structure remains intact.

**Figure 3.**
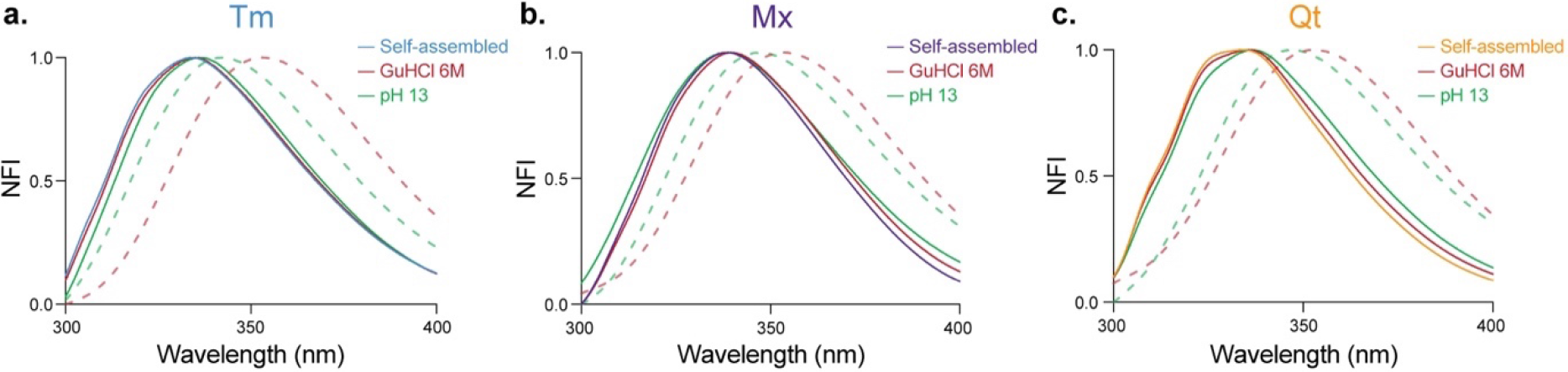
Comparison of Enc spectra of self-assembled, disassembled in GuHCl 6 M or pH 13, and subsequent reassembly. **(a)** *Tm-Enc* ITF emission when self-assembled (blue), disassembled with 6 M GuHCl (dotted red), reassembled after 6 M GuHCl disassembly (solid red), disassembled with pH 13 (dotted green), and reassembled after pH 13 disassembly (solid green). **(b)** *Mx-Enc* ITF emission when self-assembled (purple), disassembled with 6 M GuHCl (dotted red), reassembled after 6 M GuHCl disassembly (solid red), disassembled with pH 13 (dotted green), and reassembled after pH 13 disassembly (solid green). **(c)** *Qt-Enc* ITF emission when self-assembled (orange), disassembled with 6 M GuHCl (dotted red), reassembled after 6 M GuHCl disassembly (solid red), disassembled with pH 13 (dotted green), and reassembled after pH 13 disassembly (solid green).

Reassembly of all encapsulins after disassembly in either 6 M GuHCl or pH 13 was initiated via overnight dialysis into reassembly buffer. Following disassembly in 6 M GuHCl, *Tm-Enc, Mx-Enc,* and *Qt-Enc* all reassembled back to their original structures, as indicated by ITF maximum emission and DLS size (Figure 3a–c, Supplementary Figure S3). In contrast, after disassembly in pH 13, only *Tm-Enc* reassembled to its original structure. For *Mx-Enc* and *Qt-Enc*, a blue-shift in ITF emission, resembling the Trp maximum emission of the original self-assembled material, was observed. However, native-PAGE results showed no bands (Supplementary Figure S3), and TEM analysis found no structures for *Mx-Enc,* and only a single nanocage was imaged for *Qt-Enc* (Figure 2i). Indeed, DLS analysis of pH 13 reassembled *Mx-Enc* and *Qt-Enc* showed the presence of either large aggregates or smaller structures (~17 nm), which suggests the subunits may be reassembling into smaller cages and/or intermediate structures at a concentration too low to detect by native-PAGE and TEM.

As previously discussed, at pH 13, hydrolysis of the peptide bonds can occur, leading to protein misfolding and/or aggregation^44^, which may account for some of this inefficiency in encapsulin reassembly. Interestingly, although the position of the Trp emission peak for the original self-assembled, GuHCl reassembled and pH 13 reassembled samples of *Qt-Enc* are not significantly different, a difference in spectral shape is noted for the pH 13 reassembled sample. The blue-shifted shoulder of the *Qt-Enc* Trp emission peak, thought to correspond to Trp95 buried in the pentamer/hexamer interface of the intact nanocage, is lower in intensity for the pH 13 reassembled sample. This result is consistent with reassembly of a different structure with a native-like fold, instead of the original intact *T* = 4 *Qt-Enc* nanocage, upon reassembly from pH 13. The reassembly of *Mx-Enc* and *Qt-Enc* after pH 13 disassembly into smaller nanocages and/or alternative structures in inefficient quantities suggests this pathway, in its native form, may not be suitable for some applications.

### Timescale of encapsulin reassembly

To advance encapsulins as a cargo-carrying platform, disassembled encapsulin needs to be able to be reassembled on a viable timescale. The encapsulin nanocages were disassembled for 1 h via 6 M GuHCl or pH 13 before reassembly was initiated using dialysis, with samples measured with ITF and DLS every 15 min over a 75 min timeframe. The rate of reassembly appeared to vary between each Enc. *Tm-Enc* reassembled faster than *Mx-Enc* and *Qt-Enc*, with DLS indicating *Tm-Enc* reached its assembled diameter within 15 min after being disassembled via pH 13, and between 15–30 min when disassembled via 6 M GuHCl (Figure 4a & b, Supplementary Table S5 & S6). The faster reassembly after pH 13 disassembly compared to GuHCl may be explained by the subunits still maintaining some structure after incubation in pH 13 conditions. However, *Tm-Enc* reassembly after pH 13 disassembly displayed a more gradual Trp burial and a slight decrease in size at 45 mins (Figure 4a & b, Supplementary Table S6). This suggests reassembly after pH 13 may follow a more dynamic pathway, with rapid nanocage formation, followed by restructuring. In contrast, after 6 M GuHCl disassembly, *Qt-Enc* reassembled into its original diameter between 30–45 min, and *Mx-Enc* appeared to partially assemble between 45–60 min, but only completely assembled between 60-75 min (Figure 4a & b, Supplementary Table S5). The difference in rates between encapsulins may reflect the overall stability of the complex i.e., complexes that are more stable may reassemble quicker. In a recent report, the *in vivo* loading of cargo proteins into *Tm-Enc* during its self-assembly was found to be ~8 times less efficient than with *Mx-Enc*.^47^ The authors suggested that this striking difference was partly due to the smaller *Tm-Enc* self-assembling at a faster rate than *Mx-Enc*, thus limiting the available contact time between ESig-tagged cargo and the nanocage’s interior surface. This hypothesis is now supported by our observation that *Tm-Enc* reassembles up to four times quicker than the larger *Mx-Enc* (Figure 4). This therefore highlights the critical role assembly time-frames play in cargo-loading efficiency, and that modifying such properties can enhance cargo packing, density and/or stoichiometry.

**Figure 4.**
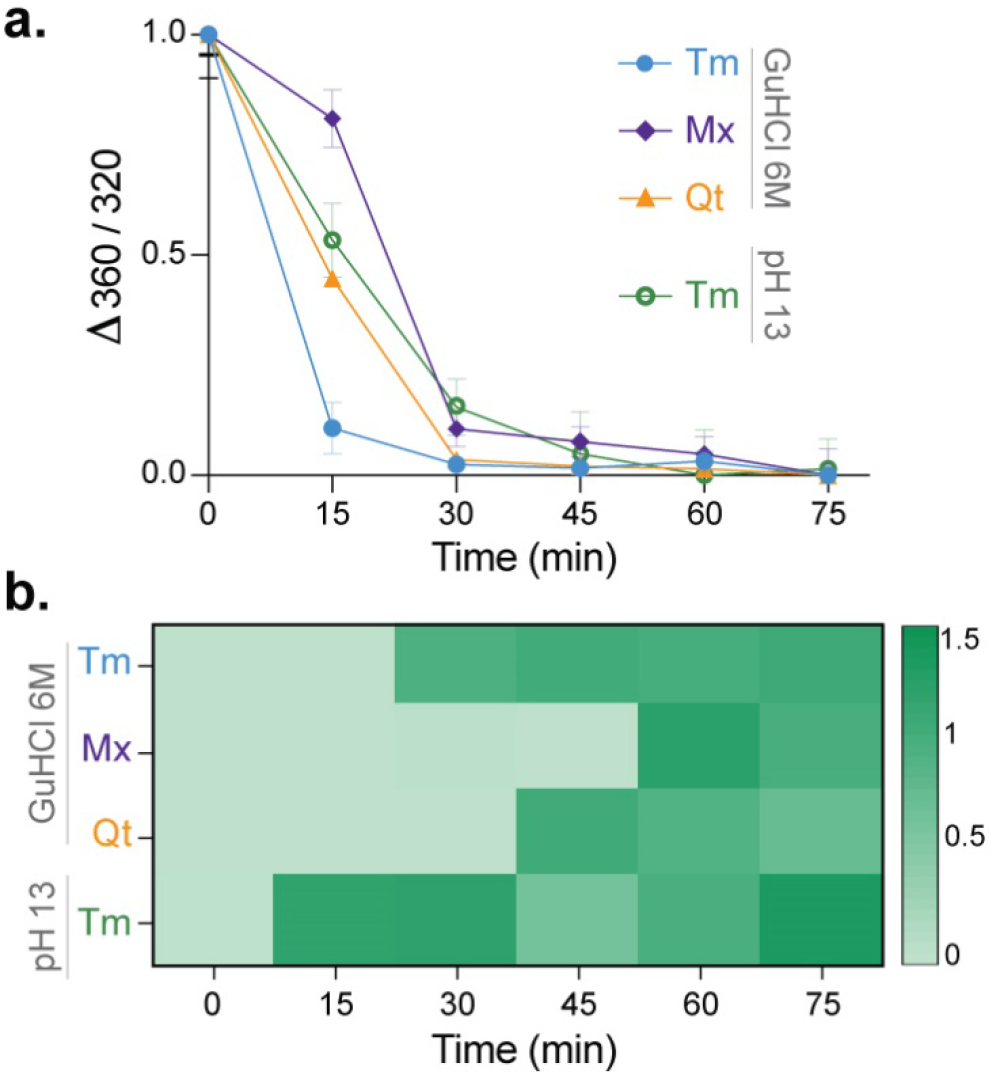
Reassembly timeframes of encapsulins. **(a)** ITF shift in emission wavelength (Δ360/320) of *Tm-Enc*, *Mx-Enc*, and *Qt-Enc* disassembled via 6 M GuHCl and *Tm-Enc* disassembled via pH 13 and reassembly measured every 15 min. The difference in emission wavelength of disassembled Enc (normalised to 1) and assembled Enc at 75 min (normalised to 0) was plotted. Error bars represent the mean ± standard deviation; n = 3 from three independent experiments. **(b)** Heatmap of DLS measurements of *Tm-Enc, Mx-Enc,* and *Qt-Enc* disassembled via 6 M GuHCl and *Tm-Enc* disassembled via pH 13 and reassembly measured every 15 min. DLS results were normalised so that 1 = expected assembled size and 0 = disassembled encapsulin.

The rapid burial of Trp residues for all encapsulins within 15 min, as shown by ITF, prior to formation of the cages, suggests a reassembly pathway where subunits first fold into an intermediate structure before gradually forming the cage. A previous study on encapsulin from *B. linens* (*T* = 1) suggested reassembly occurred with subunits first forming stable dimers prior to the final nanocage formation, with a preference for even numbered stoichiometries as demonstrated via mass spectrometry analysis.^38^ Additionally, ferritin has been found to have a biphasic reassembly, where an initial fast step occurs with folding of subunits and unknown stable intermediates, followed by the slower restructuring of intermediates into the nanocage.^23^ Data of pH 13 reassembly rate for *Mx-Enc* and *Qt-Enc* was not included, as the presence of aggregation prevented extraction of accurate DLS size values. This suggests that reassembly from the pH 13 disassembled state for the larger and more complex *Mx-Enc* and *Qt-Enc* may be prone to protein misfolding pathways and/or partial assembly, and thus, without additional re-engineering, may not be ideal for use in future applications.

### Nanomechanical stability of self-assembled vs reassembled encapsulins

To understand any changes in the structural integrity of *Tm-Enc, Mx-Enc,* and *Qt-Enc* after reassembly in the conditions tested above, AFM was utilised to compare the morphology, rupture point, and elasticity between self-assembled and reassembled encapsulins. Previous AFM analysis has been done on the rigidity of *Tm-Enc* and encapsulin from *B. linens* (*T* = 1), and suggested that the presence of cargo within encapsulin from *B. linens* may lead to some destabilisation, as indicated by a lower rupture force in loaded versus unloaded encapsulin.^38^ However, the effect of reassembly on the nanomechanical stability on *Tm-Enc,* as well as the larger and more structurally complex *Mx-Enc* and *Qt-Enc,* is yet to be elucidated.

Individual encapsulins were imaged with a scan size of ~100 nm × 100 nm to visualise their topographical detail. Self-assembled encapsulins were found to be at their expected size, including both *T* = 3 and *T* = 1 conformations of *Mx-Enc* (Figure 5a). However, clear morphological variation can be seen after reassembly, with all encapsulins demonstrating flattening of their surfaces. Additionally, *Qt-Enc* appeared more aggregated after reassembly.

**Figure 5.**
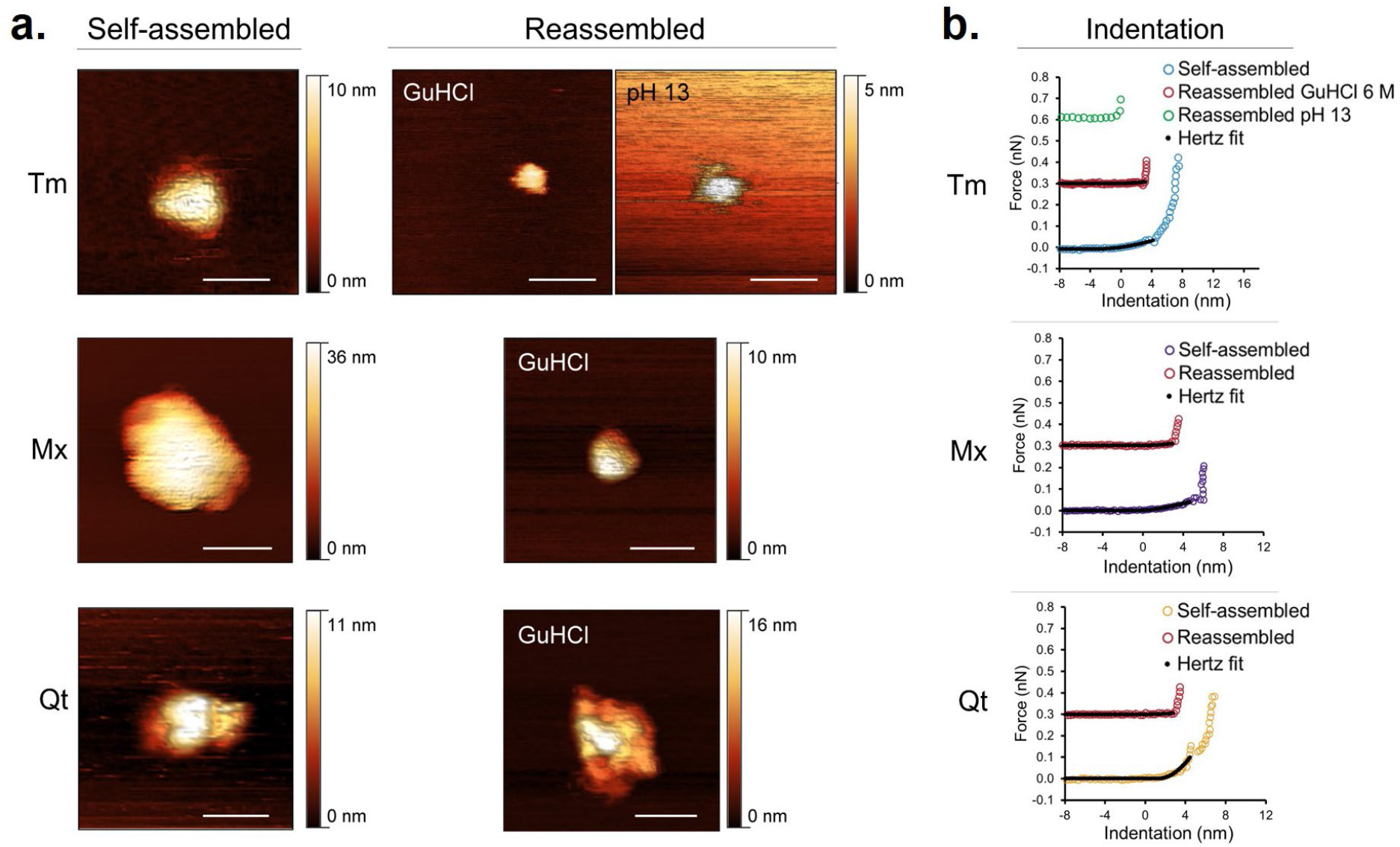
AFM imaging and elasticity comparing self-assembled/reassembled encapsulins. **(a)** Nanoscale AFM images obtained for individual encapsulins comparing morphologies of self-assembled *Tm-Enc, Mx-Enc,* and *Qt-Enc* with encapsulins disassembled via pH 13 or GuHCl (6 M) and subsequently reassembled. White scale bars represent 30 nm. **(b)** Representative indentation curves and Hertz Fit of encapsulins used to determine Young’s elastic modulus (E) using Hertz equation to convert experimentally derived force versus distance curves into indentation data (coloured circles) and to fit the elastic (non-linear) region to calculated E values (black lines).

Being composed of similar constituents, approximately the same elasticity would be expected between each self-assembled encapsulin, which was found to be the case of *Tm-Enc* and *Mx-Enc,* with an average elasticity of 2.77 ± 0.96 and 2.12 ± 0.81 MPa, respectively (Figure 5b, Supplementary Figure S5). However, *Qt-Enc* displayed a higher average elasticity of 25.60 ± 14.92 MPa, indicating it has greater structural integrity and is therefore more resilient to deformation (Figure 5b, Supplementary Figure S5). *Qt-Enc* may therefore be more strongly self-assembled than *Tm-Enc* and *Mx-Enc,* which could be attributed to its higher structural complexity, as the different symmetries and subunit-subunit contacts may influence the strength of interfacial interactions.^48^ Upon reassembly following 6 M GuHCl disassembly, the elasticity of all encapsulins decreased significantly, with *Tm-Enc* average elasticity 1.16 ± 0.93 MPa (58.12% decrease), *Mx-Enc* 1.39 ± 0.78 MPa (34.43% decrease), and *Qt-Enc* 0.85 ± 0.62 MPa (96.68% decrease), indicating a significant reduction in structural integrity (Figure 5b, Supplementary Figure S5). Elasticity data of reassembled *Tm-Enc* after disassembly via pH 13 was not included, as, during testing, the sample became unstructured and did not retain any elasticity or show any elastic response. This suggests that even though *Tm-Enc* reassembled after disassembly via pH 13, these conditions may have irreversibly altered the protein by hydrolysis, thereby affecting stability.^44^

The rupture force represents the maximum force a protein cage withstands prior to puncture by the AFM tip.^49^ The rupture force of all self-assembled encapsulins were between ~0.1-0.2 nN, indicating they are fairly fragile (Figure 5b). This is in contrast to AFM results from a previous study on self-assembled empty *Tm-Enc* and encapsulin from *B. linens,* which had rupture forces of 0.63 and 0.64 nN, respectively^38^, indicating that the encapsulins used in the current study are slightly more fragile. After reassembly, the rupture force decreased further by a factor of ~5 for each encapsulin system tested (Figure 5b).

Together, these results show that, after undergoing disassembly in either 6 M GuHCl or pH 13 and subsequent reassembly, all three encapsulin architectures significantly lose structural integrity. In terms of encapsulins loaded with cargo under *in vitro* conditions, the observed increased fragility could be beneficial if wanting a system that can be easily broken down to release cargo, such as intracellular drug delivery or immunotherapy. If in need of a more robust system (e.g., enzyme-loaded nanocages for industrial biocatalysis), however, these results demonstrate the importance to understand and modify encapsulins so they can disassemble in more benign conditions. Furthermore, to enhance understanding of the reassembly pathway of encapsulins, future studies could assess real-time assembly of the subunits. This has been achieved with large bacterial microcompartments, where AFM revealed that the shell facets assemble from preformed oligomers.^50^

## Conclusion

In summary, this study characterised the conditions facilitating the disassembly and reassembly of *Tm-Enc, Mx-Enc,* and *Qt-Enc*, with each encapsulin system varying in degree of sensitivity to GuHCl, pH, and temperature. When disassembled in high pH conditions, the subunits appeared to individually remain structurally intact, as opposed to the complete unravelling observed when disassembled in high concentrations of GuHCl. Furthermore, *Tm-Enc, Mx-Enc,* and *Qt-Enc* were all able to reassemble within 75 min after disassembly with GuHCl; however, only *Tm-Enc* was able to reassemble after disassembly at pH 13 (within 15 min). Our results suggest that disassembly by GuHCl and pH 13 affect different interfacial interactions. Self-assembly, disassembly, and reassembly of protein nanocages are highly dependent on interfacial subunit interaction^17^, and therefore, further understanding of these interactions is a fundamental step required to harness their potential. Furthermore, using AFM, we revealed a previously unknown effect of *in vitro* disassembly/reassembly on encapsulin stability, finding that encapsulin loses a substantial amount of structural integrity when disassembled and subsequently reassembled. These findings have implications if requiring *in vitro* loaded encapsulin for a particular application, but may be beneficial when nanocage dissociation and cargo release is favored.

In high concentrations of GuHCl, all encapsulins completely unfold and refold efficiently; however, these harsh conditions pose limitations for applications involving *in vitro* loading of cargo that may be sensitive to these conditions, as it could be destroyed in the process. Furthermore, the need to use high concentrations of GuHCl or alkaline pH in order to achieve disassembly may limit use for biomedical applications, where biocompatibility is paramount. Therefore, understanding the disassembly/reassembly conditions is an important step to elucidate target regions within encapsulin subunits that can be engineered to allow nanocage disassembly to occur under milder conditions. For example, the elucidation of ferritin disassembly/reassembly conditions has allowed for targeted amino acid engineering to enhance its potential as a nanocarrier for pH-sensitive compounds. As previously discussed, ferritin requires strong acidic conditions (pH ≤ 2) to disassemble, and molecules sensitive to these conditions are unable to be loaded. Therefore, by removing the last 23 amino acids from the carboxyl terminal, researchers found that ferritin was able to disassemble at pH 4 and reassemble again at pH 7, with the ability to load the bioactive agent curcumin and maintain its regular structure.^24^

Engineering interfaces can also allow enhanced control over size and cargo-loading efficiency of protein nanocages. For instance, due to being disassembled under high acidic conditions, a lot of ferritin is lost during this process and therefore it has a low loading efficiency. Consequently, ferritin variant subunit interfaces were engineered to display His6 motifs, which allowed nanocages to reassemble in the presence of either transition metal ions or pH 10, which increased loading efficiency by 1.6-3.6 times compared with previous acid disassembly.^51^ Due to ferritin being limited to one size, researchers engineered the subunit interface by first identifying the key subunit interfaces and removing the amino acids that do not participate in the interfacial interactions, resulting in the formation of a 17 nm 48-er nanocage instead of the 12 nm 24-er native ferritin structure.^52^ With an increased understanding of encapsulin structure, disassembly/reassembly conditions, molecular mechanisms, stability, and stoichiometry of intermediate structures, future genetic engineering efforts of subunits may similarly lead to enhanced control over conditions that trigger disassembly/reassembly, size, structure, and cargo-loading efficiency of encapsulins.

In summary, the findings of this study advance our understanding of encapsulins by providing critical insight into their unique disassembly/reassembly dynamics. This knowledge provides a roadmap towards an encapsulin “tool-kit” comprised of nanocages with varying structural architectures and biochemical/biophysical properties, that can be readily selected and further customised for a specific nanobiotechnological application.

## ASSOCIATED CONTENT

### Supporting Information

List of genetic constructs, examples of purification chromatographs, gel images of purified encapsulins, molecular graphics of Trp residue positions within encapsulin subunits, gel images of disassembled/reassembled encapsulins, stability study of encapsulins in REDOX and ionic condition, AFM elasticity histograms, and DLS measurement tables.

## AUTHOR INFORMATION

Corresponding Author Andrew Care

## Author Contributions

I.B co-designed the research, generated all nanocage constructs, conducted all disassembly/reassembly characterisation work, performed data analysis and wrote the manuscript. S.G assisted in experimental design, data interpretation, and wrote the manuscript. D.D assisted in protein production and purification, TEM imaging, protein modelling, and revised the manuscript. A.E performed AFM experiments and data analysis. L.C.P supervised the project and revised the manuscript. A.C. conceptualised and co-designed the study, supervised the project, and wrote the manuscript.

## Notes

The authors declare no competing financial interest.

## ACKNOWLEDGMENTS

I.B. is supported by a Dementia Australia Research Foundation PhD scholarship. A.C. is supported by the UTS Chancellor’s Postdoctoral Research Fellowship scheme. This work was supported by grants to A.C and L.C.P by the Dementia Australia Research Foundation, the National Foundation for Medical Research and Innovation, the Brain Foundation and the NeuroSurgical Research Foundation.

