## Supplementary Material for "Exploring the Self-Assembly of Encapsulin Protein Nanocages from Different Structural Classes"

**Supplementary Table S1.** Expression plasmids constructed for this study.

| Plasmids | Description | Protein ID | Selection antibiotic |
|---|---|---|---|
| pETDuet-1_Tm | Encapsulin from *Thermotoga maritima (Tm)* | UniProt: Q9WZP3 | Carbenicillin |
| pACYC-Duet-1_Mx | Encapsulin from *Myxococcus xanthus (Mx)* | UniProt: Q1D6H4 | Chloramphenicol |
| pACYC-Duet-1_Qt | Encapsulin from *Quasibacillus thermotolerans (Qt)* | UniProt: A0A0F5HPP7 | Chloramphenicol |

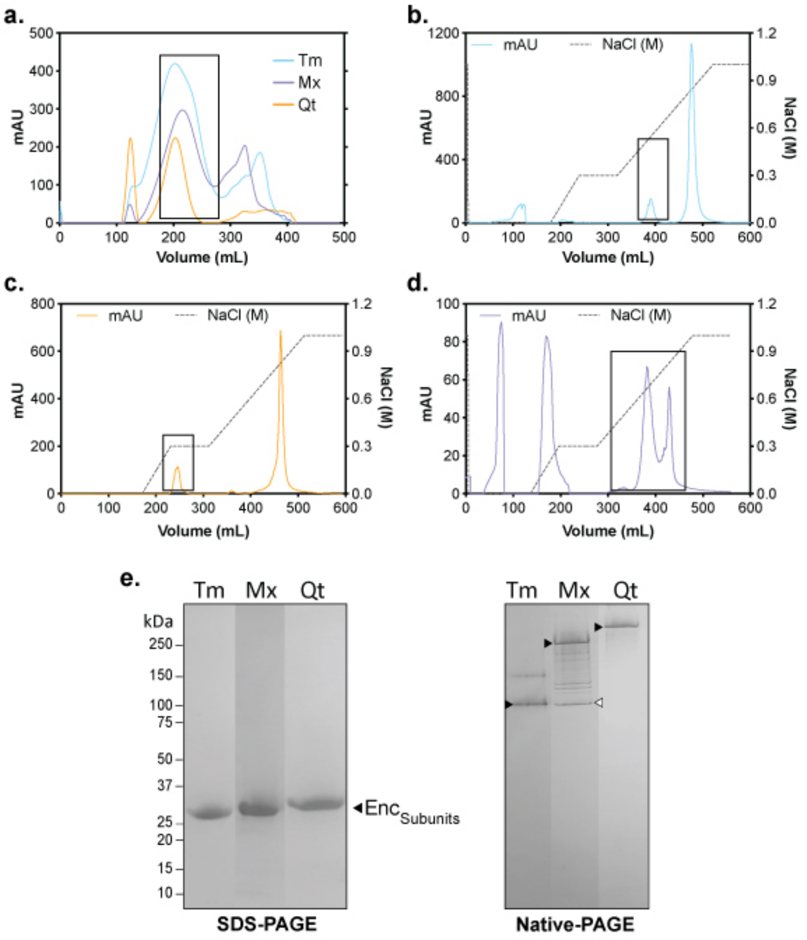


**Supplementary Figure S1. Example chromatograms of encapsulin (Enc) nanocage purifications. (a)** Size exclusion chromatography (SEC) of *Tm-Enc* (blue), *Mx-Enc* (purple) and *Qt-Enc* (orange) nanocages. The proteins of interest (black square) eluted between 170-280 mL. Anion exchange chromatography of SEC-purified **(b)** *Tm-Enc*, **(c)** *Mx-Enc*, and **(d)** *Qt-Enc*. Peaks that correspond to the proteins of interest are highlighted within a black square. **(e)** Coomassie-stained SDS-PAGE showing the purification of $Enc_{Subunit}$ *Tm-Enc* 30.5 KDa, *Mx-Enc* 31.6 KDa and *Qt-Enc* 32.2 KDa. Blue-Native PAGE showing the purification of assembled *Tm-Enc* (black triangle *T* = 1), *Mx-Enc* (black triangle *T* = 3, white triangle *T* = 1) and *Qt-Enc* (black triangle *T* = 4).

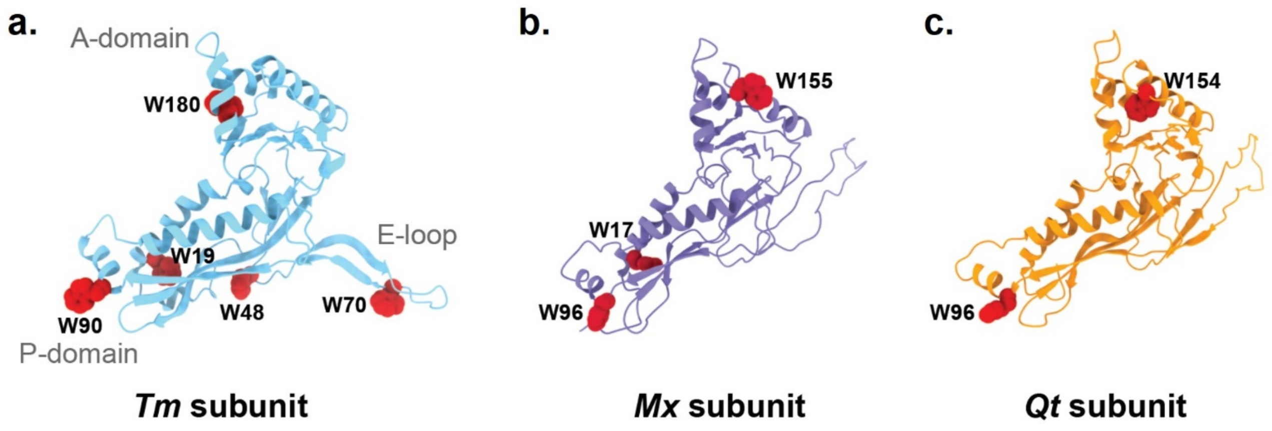


**Supplementary Figure S2**. Structures of the individual subunits from **(a)** *Tm-Enc* (PDB: 3DKT), **(b)** *Mx-Enc* (PDB: 4PT2), and **(c)** *Qt-Enc* (PDB: 6NJ8). All tryptophan (W) residues are highlighted in red: ***Tm***: W19, W48, W70, W90 and W180; ***Mx***: W17, W96 and W155; ***Qt***: W95 and W154. Molecular graphics were created using UCSF ChimeraX.

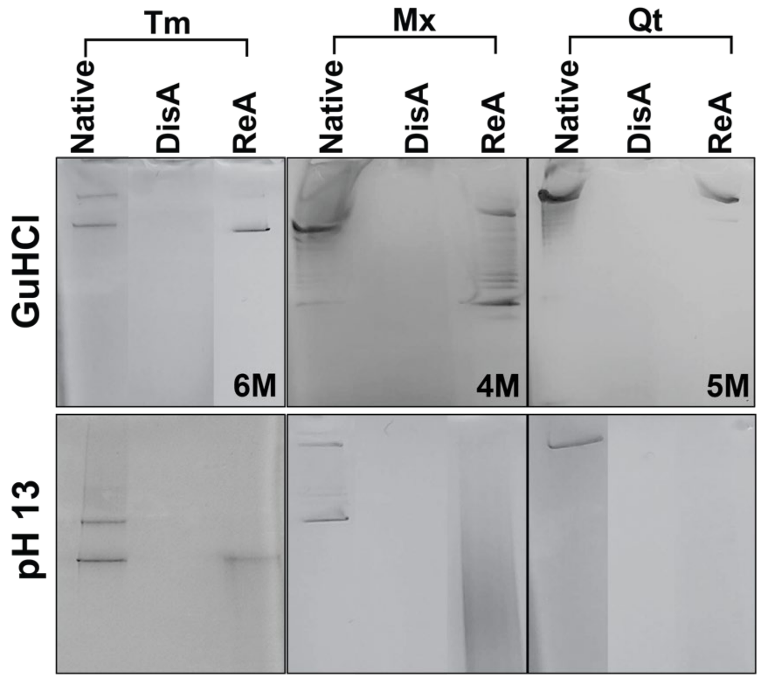


**Supplementary Figure S3**. pH or guanidine hydrochloride (GuHCl) triggered disassembly/ reassembly of encapsulins. Blue Native-PAGE showing purified *Tm-Enc, Mx-Enc* and *Qt-Enc* proteins self-assembled (native), disassembled (DisA) with GuHCl (upper panel) or pH13 (bottom panel) and reassembled (ReA) after pH or denaturant treatment.

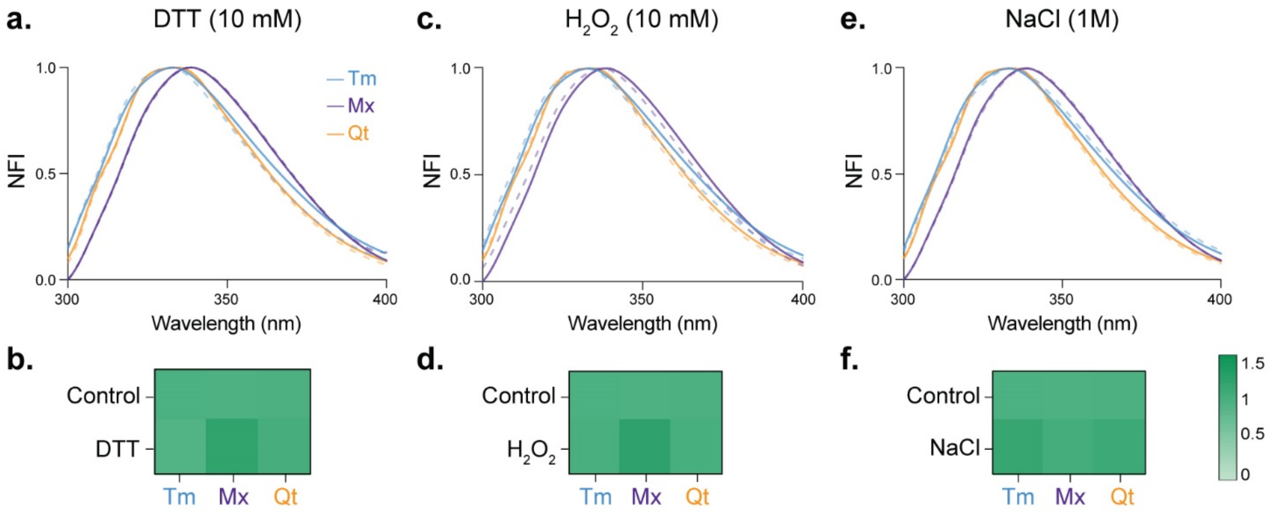


**Supplementary Figure S4. Stability of Encs in REDOX and ionic conditions. (a)** ITF emission peak of Encs incubated in 10 mM DTT compared to controls. **(b)** Heat map of DLS of Encs incubated in 10 mM DTT. **(c)** ITF emission peak of Encs incubated in 10 mM $H_2O_2$ compared to controls. **(d)** Heat map of DLS of Encs incubated in 10 mM $H_2O_2$. **(e)** ITF emission peak of Encs incubated in 1 M NaCl compared to controls. **(f)** Heat map of DLS of Encs incubated in 1 M NaCl. Solid lines represent controls, and dotted lines represent Encs incubated in DTT, $H_2O_2$, or NaCl. DLS results were normalised so that 1 = expected assembled size and 0 = disassembled encapsulin.

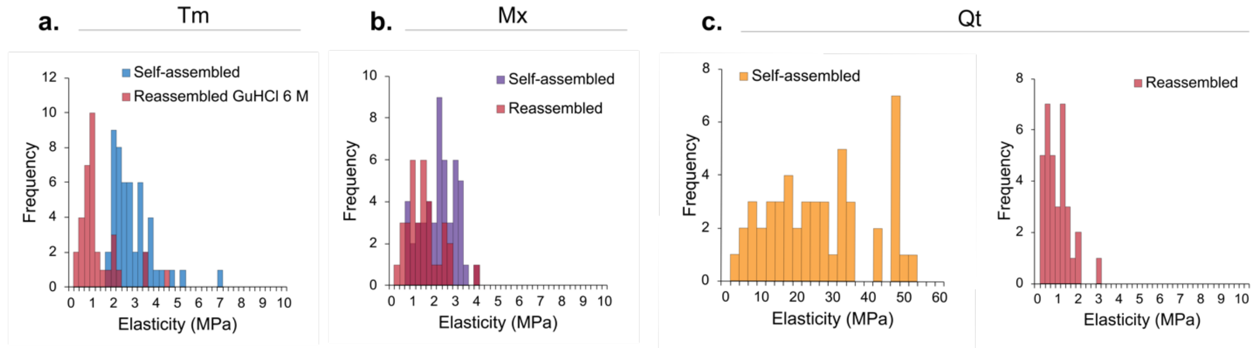


**Supplementary Figure S5. AFM elasticity comparing self-assembled/reassembled encapsulins.** The distribution of Young's modulus values for self-assembled and reassembled **(a)** *Tm-Enc*, **(b)** *Mx-Enc*, and **(c)** *Qt-Enc*.

**Supplementary Table S2.** DLS measurements after encapsulins' disassembly at different concentrations of guanidine hydrochloride (GuHCl)

| GuHCl (M) | Tm | | Mx | | Qt | |
|---|---|---|---|---|---|---|
| | Size by number (nm) | St dev (nm) | Size by number (nm) | St dev (nm) | Size by number (nm) | St dev (nm) |
| 0 | 33.21 | 9.29 | 32.05 | 5.47 | 38.99 | 7.01 |
| 1 | 25.22 | 8.96 | 33.52 | 7.53 | 37.89 | 7.48 |
| 2 | 19.71 | 2.88 | 31.13 | 6.38 | 38.60 | 7.47 |
| 3 | 37.93 | 6.10 | <0.5 | 0.07 | 50.82 | 12.40 |
| 4 | <0.5 | 0.08 | <0.5 | 0.05 | <0.5 | 0.05 |
| 5 | <0.5 | 0.07 | <0.5 | 0.05 | <0.5 | 0.07 |
| 6 | <0.5 | 0.04 | <0.5 | 0.07 | <0.5 | 0.03 |
| 7 | <0.5 | 0.06 | <0.5 | 0.06 | <0.5 | 0.04 |

**Supplementary Table S3.** DLS measurements after encapsulins' disassembly at different pH

| pH | Tm | | Mx | | Qt | |
|---|---|---|---|---|---|---|
| | Size by number (nm) | St dev (nm) | Size by number (nm) | St dev (nm) | Size by number (nm) | St dev (nm) |
| 3 | 23.36 | 5.33 | 30.88 | 4.01 | 35.40 | 2.40 |
| 4 | 24.46 | 5.92 | 30.90 | 6.67 | 34.94 | 8.10 |
| 5 | 26.03 | 6.43 | 33.76 | 4.79 | 38.70 | 5.28 |
| 6 | 23.98 | 4.89 | 30.22 | 5.74 | 40.70 | 5.38 |
| 7 | 24.56 | 5.12 | 30.92 | 5.95 | 41.45 | 5.82 |
| 8 | 25.60 | 5.47 | 33.32 | 4.78 | 42.21 | 4.96 |
| 9 | 25.06 | 5.48 | 31.60 | 5.79 | 41.30 | 5.45 |
| 10 | 22.76 | 5.46 | 31.39 | 6.00 | 38.89 | 6.49 |
| 11 | 24.95 | 5.39 | 30.85 | 5.81 | 40.96 | 5.63 |
| 12 | 28.49 | 3.75 | 13.17 | 1.04 | 40.74 | 3.53 |
| 13 | <0.7 | 0.04 | <0.7 | 0.04 | <0.7 | 0.12 |

**Supplementary Table S4.** DLS measurements after encapsulins' stability at different temperatures

| Temperature (°C) | TM | | MX | | QT | |
|---|---|---|---|---|---|---|
| | Size by number (nm) | St dev (nm) | Size by number (nm) | St dev (nm) | Size by number (nm) | St dev (nm) |
| 20 | 23.99 | 5.08 | 31.79 | 5.06 | 39.28 | 6.87 |
| 30 | 22.71 | 4.98 | 27.32 | 8.14 | 38.79 | 7.06 |
| 40 | 22.77 | 5.05 | 32.02 | 5.16 | 39.43 | 6.85 |
| 50 | 23.05 | 5.25 | 31.57 | 5.04 | 39.39 | 6.56 |
| 60 | 22.44 | 4.76 | 33.14 | 3.67 | 39.87 | 5.40 |
| 70 | 22.68 | 4.63 | 31.83 | 6.41 | 36.84 | 5.36 |
| 80 | 23.26 | 4.56 | <0.5 | 0.02 | <0.5* | 0.03 |
| 90 | 24.16 | 4.22 | <0.5 | 0.02 | <0.5** | 0.07 |
| Cooled back to 20 | 20.34 | 4.43 | 784.2 | 87.26 | 34.11 | 7.70 |

*33.1% of population was ~38.85 nm. **16% of population was ~13 nm.

**Supplementary Table S5.** DLS measurements over time of encapsulins' reassembly after disassembly with 6M guanidine hydrochloride

| Time (min) | Tm | | Qt | | Mx | |
|---|---|---|---|---|---|---|
| | Size by number (nm) | St dev (nm) | Size by number (nm) | St dev (nm) | Size by number (nm) | St dev (nm) |
| 0 | <0.5 | 0.05 | <0.5 | 0.06 | <0.5 | 0.06 |
| 15 | <0.5 | 0.05 | <0.5 | 0.09 | <0.5 | 0.07 |
| 30 | 24.24 | 4.86 | <0.5 | 6.73 | 2.00 | 0.13 |
| 45 | 26.62 | 6.16 | 42.19 | 10.76 | 0.70 | 0.13 |
| 60 | 25.73 | 5.17 | 36.68 | 9.26 | 10.75 | 1.22 |
| 75 | 26.96 | 4.54 | 30.25 | 8.65 | 33.54 | 7.06 |

**Supplementary Table S6.** DLS measurements over time of encapsulins' reassembly after disassembly at pH 13

| Time (min) | TM | |
|---|---|---|
| | Size by number (nm) | St dev (nm) |
| 0 | <0.8 | 0.08 |
| 15 | 30.69 | 1.96 |
| 30 | 30.79 | 5.91 |
| 45 | 16.11 | 1.92 |
| 60 | 24.72 | 4.47 |
| 75 | 34.78 | 5.85 |

**Supplementary Table S7.** DLS measurement of encapsulins' stability in REDOX or ionic conditions.

| Encapsulin | Treatment | Size by number (nm) | St dev (nm) |
|---|---|---|---|
| Tm | Control | 23.70 | 4.90 |
| | 10 mM $H_2O_2$ | 24.70 | 7.30 |
| | 10 mM DTT | 23.00 | 6.40 |
| | 1M NaCl | 29.20 | 8.50 |
| Mx | Control | 32.00 | 6.00 |
| | 10 mM $H_2O_2$ | 41.30 | 6.80 |
| | 10 mM DTT | 40.80 | 6.90 |
| | 1M NaCl | 34.55 | 7.10 |
| Qt | Control | 38.10 | 7.30 |
| | 10 mM $H_2O_2$ | 39.40 | 7.00 |
| | 10 mM DTT | 40.40 | 7.70 |
| | 1M NaCl | 43.56 | 9.94 |
